# *In vivo* epigenetic editing of *sema6a* promoter reverses impaired transcallosal connectivity caused by *C11orf46/ARL14EP* neurodevelopmental risk gene

**DOI:** 10.1101/491779

**Authors:** Cyril J. Peter, Atsushi Saito, Yuto Hasegawa, Yuya Tanaka, Gabriel Perez, Emily Alway, Sergio Espeso-gil, Tariq Fayyad, Chana Ratner, Aslihan Dincer, Achla Gupta, Lakshmi Devi, John G. Pappas, François M. Lalonde, John A. Butman, Joan C. Han, Schahram Akbarian, Atsushi Kamiya

## Abstract

Many neuropsychiatric risk genes contribute to epigenetic regulation of gene expression but very little is known about specific chromatin-associated mechanisms governing the formation and maintenance of neuronal connectivity. Here we show that transcallosal connectivity is critically dependent on *C11orf46* (also known as *ARL14EP*), a small nuclear protein encoded in the chromosome 11p13 *Wilms Tumor, Aniridia, Genitourinary Abnormalities, intellectual disability (formerly referred to as Mental Retardation)* (WAGR) risk locus. *C11orf46* haploinsufficiency in WAGR microdeletion cases was associated with severe hypoplasia of the corpus callosum. *In utero* short hairpin RNA-mediated C11orf46 knockdown disrupted transcallosal projections of cortical pyramidal neurons, a phenotype that was rescued by wild type C11orf46 but not the C11orf46^R236H^ mutant associated with autosomal recessive intellectual disability. Multiple genes encoding key regulators of axonal growth and differentiation, including *Sema6A*, were hyperexpressed in C11orf46-knockdown neurons. Importantly, RNA-guided epigenetic editing of neuronal *Sema6a* gene promoters via a dCas9 protein-conjugated SunTag scaffold with multimeric (10x) C11orf46 binding during early developmental periods, resulted in normalization of expression and rescue of transcallosal dysconnectivity via repressive chromatin remodeling, including up-regulated histone H3K9 methylation by the KAP1-SETDB1 repressor complex. Our study demonstrates that interhemispheric communication is highly sensitive to locus-specific remodeling of neuronal chromatin, revealing the therapeutic potential for shaping the brain’s connectome via gene-targeted designer activators and repressor proteins.

## Introduction

A wide range of neurodevelopmental disorders manifesting in infancy and early childhood (including intellectual disability and autism spectrum disorder) or young adulthood (including schizophrenia) are associated with disrupted interhemispheric communication, which in some but not all of the affected cases is accompanied by structural alterations of the corpus callosum, the brain’s largest commissure ^1–3^. The formation of interhemispheric connectivity involves a highly orchestrated multi-step process, including midline zipper glia promoting hemispheric fusion and midline crossing of pioneer fibers, followed by ingrowth of large numbers of transcallosal axons interconnecting the left and right cerebral cortex ^4^ Early occurring disruptions of commissural development could manifest as agenesis of the corpus callosum ^4^ However, alterations affecting later phases of development, including defective or misdirected axonal growth cone navigation with failure to cross the midline, or faulty axonal ingrowth into the contralateral cortex, could be responsible for the partial thinning, or developmental hypoplasia of the corpus callosum, extending either across its entire rostro-caudal axis or subportion thereof ^4,5^. In addition, proper region-and lamina-specific axonal innervation and arborization within contralateral cortex are dependent on the functional activity of cortical projection neurons ^6^. Furthermore, late occurring defects of transcallosal connectivity may not result in overt macroscopic defects of brain morphology including its commissures. Such a complex developmental program of commissural connectivity is, perhaps unsurprisingly, associated with a heavy footprint in the genetic risk architecture of neurodevelopmental disease. In fact, a number of chromosomal copy number variants (CNV) have been linked to callosal agenesis ^7^.

Of note, many cases with a chr. 22q11.2 micro-deletion or -duplication ^8^ as one of the most frequently diagnosed CNV in neuropsychiatric disease cohorts are affected by callosal hypo- and hyperplasia ^8^ and furthermore, interhemispheric dysconnectivity phenotypes have been associated with a rapidly increasing list of specific point mutations in cell surface signaling genes encoding key regulators of neurite outgrowth and connectivity, such as the axon guidance receptor *ROBO1 ^9^* or the *L1CAM* adhesion molecule ^10^. Much less is known about the role of nuclear signaling molecules for disease-relevant commissural phenotypes. For example, orderly formation of transcallosal connectivity critically depends on proper gene dosage and activity of a subset of chromatin regulators such as the transcriptional repressor switch-insensitive family member A (*SIN3A*) ^11^ and the FOXG1 transcription factor ^12^. However, molecular and cellular mechanisms linking the neuronal epigenome to interhemispheric connectivity remain poorly explored.

Here, we describe *C11orf46/ARL14EP*, a small (35kDa) nuclear protein encoded in the chr. 11p13 *Wilms Tumor, Aniridia, Genitourinary Abnormalities, intellectual disability (formerly referred to as Mental Retardation)* (WAGR) risk locus, as a novel chromatin regulator of transcallosal connectivity. Importantly, we show that C11orf46 is, as an RNA-guided Cas9 fusion protein, suitable for targeted epigenomic promoter editing to drive the expression of specific regulators of axonal development in immature transcallosal projection neurons, thereby offering a molecular tool to affect interhemispheric communication.

## Results

### C11orf46 is expressed in cortical projection neurons and critical for transcallosal connectivity

A recent study identified 50 novel genes associated with autosomal recessive forms of intellectual disability, including open reading frame *C11orf46* ^13^. Importantly, moreover, this gene is located in the chr. 11p13-14 neurodevelopmental risk locus and deleted in some cases of WAGR syndrome (Online Mendelian Inheritance of Man (OMIM) #194072), a rare copy number variant disorder with its core features caused by haploinsufficiency for the *Wilms Tumor 1 (WT1)* and *PAX6* homoebox gene ^14^ Of note, *C11orf46* is positioned in between *PAX6* and the *Brain Derived Neurotrophic Factor (BDNF, chr. 11p14.1)* gene which is included in the deleted segment in approximately half of patients with WAGR syndrome and associated with obesity and more severe cognitive and behavioral deficits ^14^ However, there is evidence that, independent of *BDNF*, additional coding genes in chr. 11p13-14 could contribute to neurodevelopmental disease. For example, exome sequencing studies in Middle Eastern pedigrees associated homozygous *C11orf46 (chr11:30,323,051-30,338,458, GRCh38/hg38)* point mutations with cognitive disease and intellectual disability ^13^. To begin the neurological exploration of *C11orf46*, we studied differences in brain morphology between WAGR microdeletion cases encompassing the *C11orf46* gene and those with less extensive chr. *11p13-14.1* segment loss.

To this end, we examined magnetic resonance T1-weighted images (MRI) in N=39 WAGR deletion cases. Indeed, *C11orf46* haploinsufficiency was associated with significant differences observed in total corpus callosum (CC) and posterior CC volumes but not for other individual portions of the corpus callosum. After adjusting for age and sex, patients with isolated *PAX6*+/− (N=12) had significantly smaller total (p<0.001) and posterior (p=0.001) CC volumes than controls (N=23), a finding consistent with the role of *PAX6* in morphologic brain development ^15,16^. However, patients with combined *PAX6/C11orf46*+/− (N=27) had an even smaller posterior CC volume compared to both patients with isolated *PAX6*+/− (p=0.003) and controls (p<0.001) (**Fig. 1a,b**). Because of the established role of *BDNF* in brain development and intellectual functioning ^17^, we also considered a potential confounding effect of *BDNF*+/−. We observed no significant differences in the *PAX6/C11orf46*+/− patients with (N=17) and without (N=10) *BDNF*+/− for total or posterior CC volumes (p=0.19 and 0.47, respectively, for unadjusted comparisons; p=0.13 and 0.37, respectively, on ANCOVA adjusting for age and sex) (**Supplementary Fig. 1a**). Lymphoblastoid cell lines from *C11orf46* haploinsufficient WAGR cases have approximately 50% lower C11orf46 protein and mRNA levels, indicating that the expression of the remaining intact allele does not undergo compensatory upregulation (**Supplementary Fig. 1b**). Therefore, loss of one *C11orf46* copy in the context of chr. *11p13-14.1* microdeletions appears to be more highly detrimental for neurodevelopment and associated with more severe hypoplasia of the callosal commissure.

**Fig. 1.**
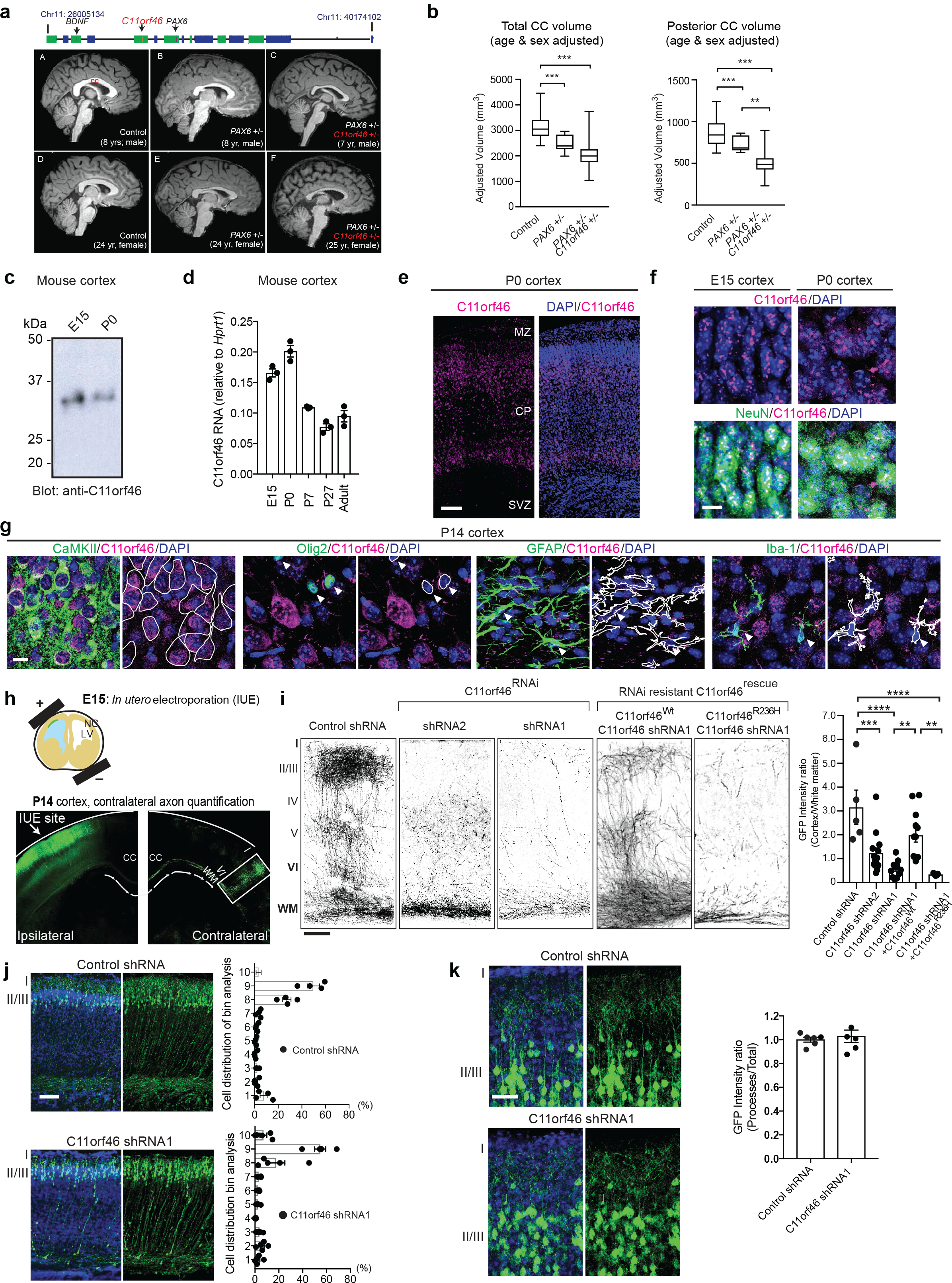
C11orf46 is a neuronal nuclear protein important for callosal development and interhemispheric connectivity. **a** (top) Genomic map of chr. 11 WAGR deletion locus. (bottom, A-F) Representative sagittal T1 MRI images 1.875 mm from midline for six subjects were shown in subpanels. Brain MRI with 1mm^3^ resolution, collected on a 3.0 T Philips Achieva MRI scanner with an 8-channel phased array head coil. Corpus callosum volumes were calculated using FreeSurfer’s (version 5.3) subcortical image processing pipeline. **A)** 8-year-old male healthy control; **B)** 8-year-old male isolated *PAX6*+/−; C) 7-year-old male heterozygous 11p13 deletion with *PAX6*+/− and *C11orf46+/−* **D)** 24-year-old female healthy control; E) 24-year-old female isolated *PAX6+/−* **F)** 25-year-old female heterozygous 11p13 deletion with *PAX6*+/− and *C11orf46*+/−. **b** Hypoplastic corpus callosum in patients with chromosome 11p13 deletion encompassing *C11orf46* is shown. ANCOVA including age and sex as covariates, compared corpus callosum volumes of healthy control (n = 23), isolated *PAX6*+/− (n = 12), and heterozygous 11p13 deletion with *PAX6*+/− and *C11orf46*+/− (n = 27). Box and whisker plots represent the distribution of corpus callosum volume from each participant. The inner bar in the box indicates median value. The upper and lower box ends represent first and third quantile, respectively. The upper and lower whisker ends represent the maximum and minimum values in the group, respectively. **c** C11orf46 protein (~35kDa) was detected by Western blotting in mouse cortices at embryonic day 15 (E15) and postnatal day 0 (P0). **d** The bar graphs represent mRNA expression of C11orf46 in the mouse cerebral cortex during prenatal (E15) and postnatal (P0, P7, P27, adult) brain development (mean± SEM). N=3. **e** C11orf46 protein (red) is predominantly expressed in the cortical plate (CP) in the developing somatosensory cortex at P0. Blue, nucleus counterstained by DAPI. Scale bar, 50μm. MZ, marginal zone; SVZ, subventricular zone. **f** C11orf46 (red) is expressed in the nucleus in punctate patterns. Majority of C11orf46 expressing cells are NeuN-positive neurons (green) at E15 and P0. Scale bar, 20μm. **g** C11orf46 is predominantly expressed in CamKII-positive pyramidal neurons, but not Olig2 (oligodendrocyte), GFAP (astrocyte), Iba-1 (microglia) -positive cells in the cerebral cortex at P14. White traces outline of cell type marker signal (Green). Scale bar, 10 p,m. **h** (top) Schematic representation of *In Utero* electroporation of GFP expressing plasmid into somatosensory cortex at E15. (bottom) P14 somatosensory cortex expressing GFP in pyramidal neurons at IUE injection site (Ipsilateral side) and axonal projections between layer I-IV (contralateral side). **i** Knockdown of C11orf46 in pyramidal neurons of layer II/III (Ipsilateral) show impaired axonal terminal arborization in the contralateral somatosensory cortex at P14. C11orf46 shRNA1 had a stronger axonal arborization impairment than shRNA2 when compared with control shRNA (first 3 panels); this axonal deficit was partially rescued by co-expressing RNAi resistant wild type C11orf46 (C11orf46^Wt^), but not by R236H mutant (C11orf46^R236H^), panels 4 and 5 from left. n = 5-14 mice per condition. *F* (4, 43)=, *P* < 0.0001. \*\**P* < 0.01, ***P < 0.001, \*\*\*\**P* < 0.0001 determined by one-way ANOVA with *post hoc* Bonferroni test. Scale bar, 100μm. **j**, **k** C11orf46 knockdown did not have an effect on radial neuronal migration at P2 (**j**) or dendritic structures of GFP-labeled pyramidal neurons in the somatosensory cortex at P14 (**k**). Scale bar, 50μm.

Having linked *C11orf46* haploinsufficiency to callosal hypoplasia, we next explored C11orf46 expression patterns in mouse brain. In the cerebral cortex, C11orf46 quantification at the level of mRNA, and protein with a custom-made anti-C11orf46 antibody (**Supplementary Fig. 1c**) showed its expression across a wide age window, with higher levels in the pre- and post-natal developmental periods as compared to the adult brain (**Fig. 1c-f**). C11orf46 was mainly expressed in the cortical plate (CP) with minimal labeling in the proliferative subventricular zone (SVZ) (**Fig. 1e**), with punctate and intranuclear distribution predominantly in neuronal nuclei as confirmed by co-staining with NeuN antibody (**Fig. 1f**). Additional double-labeling experiments confirmed that C11orf46 is predominantly expressed in Calmodulin-kinase II (CaMKII)-positive glutamatergic neurons but not in glial fibrillary acid protein (GFAP)-positive astrocytes or Olig2-positive oligodendrocytes and precursors, or Iba1-positive microglia (**Fig. 1g**). C11orf46 immunoreactivity in GABAergic neurons was much weaker than those in glutamatergic neurons (data no shown). We conclude that in the developing cerebral cortex, C11orf46 is primarily expressed in post-migratory glutamatergic cortical neurons.

To explore the role of C11orf46 in developing cortex, we knocked down its expression by *in utero* electroporation (IUE). We delivered C11orf46 (shRNA1, shRNA2) or control short hairpin RNAs (shRNAs) together with green fluorescent protein (GFP) expression plasmid using our published protocols ^18–21^. We selected embryonic day 15 (E15) for IUE procedure with two reasons: first, C11orf46 is highly expressed at E15 (**Fig. 1c,d**). Second, at this stage IUE mostly targets progenitor cells in the ventricular zone (VZ) which later differentiate into cortical layer II/III pyramidal neurons ^22,23^, approximately 80% of which become transcallosal projection neurons ^24^ as our cell type-of-interest given the clinical phenotype reported above. Indeed, E15 IUE C11orf46 knockdown resulted in severe arborization deficits in GFP-labeled callosal axons projecting into the contralateral cortex at postnatal day 14 (P14) (**Fig. 1h,i**). This phenotype was highly dependent on the level of C11orf46 knockdown. More severe axonal arborization deficits by C11orf46 shRNA1 (Fig. 1i) was associated with stronger knockdown effect on C11orf46 compared with shRNA2 (**Supplementary Fig. 1d**). Importantly, these axonal arborization deficits were partially rescued by coexpressing shRNA resistant wildtype C11orf46. Strikingly however, an shRNA resistant C11orf46 (R236H) mutant protein carrying a disease-associated non-synonymous point mutation that consists of arginine to histidine substitution at codon 236 ^13^ did not rescue the transcallosal phenotype induced by C11orf46 knockdown (**Fig. 1i** and **Supplementary Fig. 1e**). Importantly, the observed axonal phenotype resulting from C11orf46 knockdown was highly specific, because orderly radial neuronal migration into the cortex, when assessed at P2, or dendritic complexity of the transfected pyramidal neurons when assessed at P14, were indistinguishable from control (**Fig. 1j,k**).

### C11orf46 negatively regulates genes important for neurite development and axonal connectivity

Having shown that C11orf46 is widely expressed in cortical projection neurons and essential for transcallosal connectivity, we next wanted to gain initial insights into the molecular mechanisms underlying C11orf46 actions on the neuronal connectome. In the single report published to date on C11orf46 function, the protein was renamed ADP-Ribosylation Factor-Like 14 Effector Protein (ARL14EP), and ascribed with a role in major histocompatibility class (MHC) II antigen presentation in immune cells ^25^. Therefore, we were very surprised about the predominant, if not exclusive, nuclear localization of C11orf46 in our immunohistochemical assays in the developing CP (**Fig. 1e,f**). However, a recent study listed C11orf46/ARL14EP among several hundred proteins serving as nuclear export cargo for the RanGTPase-driven exportin CRM1 ^26^ and moreover, large scale total proteome protein-protein interaction mappings consistently identified chromatin regulatory proteins as C11orf46 binding partners, including the SETDB1/KMT1E (SETDB2/KMT1F)-MCAF1 histone lysine 9 (K9) methyltransferase repressor complex ^27–30^ (referred to as KMT-RC hereafter). To further examine C11orf46-associated proteins by affinity-purification coupled to mass spectrometry (**Fig. 2a**), we generated inducible HEK293 cell lines expressing Flag-tagged C11orf46 (**Fig. 2b**). We produced two types of inducible clones, one for full length FLAG-tagged C11orf46 (clones 1-5 and 10) and one for a truncated form lacking amino acids 209-237 at C11orf46’s C-terminal region which corresponds to cysteine rich domain, CRD, (C11orf46Δ) (clone 7 and 8) (**Fig. 2b** and **Supplementary Fig. 2a,b**). Interestingly, nuclear proteins outranked all other C11orf46 binding partners, with the top scoring SETDB1-KAP1-MCAF1 chromatin repressor complex ^31,32^ (**Fig. 2c-e** and **Supplementary Table 1**), which is consistent with proteome interaction databases ^27–30^. These findings were specific for full length C11orf46 because the truncated form C11orf46Δ showed only very weak, or no interactions with nuclear proteins including the SETDB1-KAP1-MCAF1 repressor complex (**Fig. 2d**). Given the *C11orf46^R236H^* point mutation in the CRD domain is required for protein binding of C11orf46 with SETDB1 (**Fig. 2b,f**), we wondered whether R236H may affect the interaction of C11orf46 and the SETDB1 complex. To test this hypothesis, in addition to inducible HEK293 cell lines expressing Flag-tagged C11orf46 and Flag-tagged C11orf46Δ, we generated cell lines for inducible expression of FLAG-C11orf46^R236H (^**Fig. 2g,h**). We observed that deletion of CRD domain dramatically weakened the interaction of C11orf46 and SETDB1, whereas the R236H substitution abolished C11orf46-SETDB1 interaction (**Fig. 2g,h**). Binding of C11orf46 with MCAF1 and KAP1, other members of the SETDB1 protein complex, were also severely decreased by R236H substitution (**Fig. 2h**). Notably, both C11orf46 with R236H mutation and that with deletion of CRD domain still kept binding affinity with histone H3 (**Fig. 2i**). Also, purified, *E.coli* derived, human C11orf46 specifically detected histone H3 tail modifications in a histone peptide array (**SupplementaryFig. 2c-e**). Similarly, reciprocal affinity purification with FLAG-tagged SETDB1 using inducible cell lines confirmed C11orf46 as a top ranking binding partner together with other members of the complex, including KAP1 and MCAF1 (**Fig. 2j-m** and **Supplementary Table 2**). Additional co-immunoprecipitation experiments with mouse anti-C11orf46 antibody in HeLa cells, and in homogenates of human cerebral cortex confirmed that C11orf46 assembles with SETDB1, MCAF1 and other regulators of repressive chromatin, including heterochromatin-associated protein 1 gamma (HP1γ) (**Fig. 2n** and **Supplementary Fig. 2f**). These results suggest that C11orf46 may function as a chromatin regulator in the SETDB1 complex (**Fig. 2o**).

**Fig. 2.**
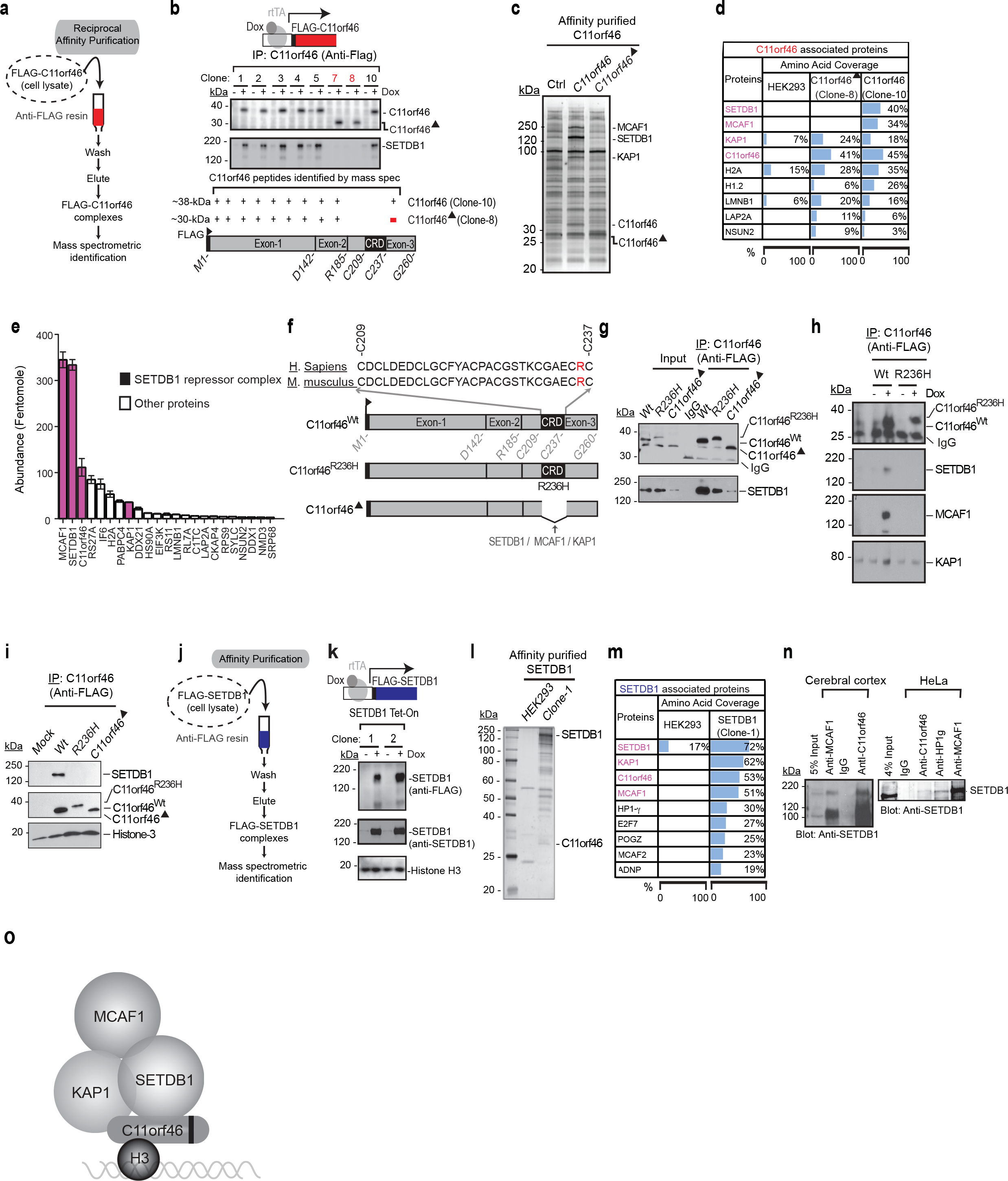
C11orf46 binds to the SETDB1 complex and regulates neuronal gene expression. **a** Flow diagrams for immunoaffinity purification of C11orf46 followed by mass spectrometric analysis. **b** Eight clones of inducible HEK293 cell lines expressing full length FLAG-tagged C11orf46 (clone 1-5 and 10) and a truncated form C11orf46 lacking amino acids 209-237 at the C-terminal region which corresponds to cysteine rich domain (CRD) of C11orf46 (C11orf46Δ) (clone 7 and 8) in the presence of doxycycline (Dox+). SETDB1 was consistently co-precipitated with full-length C11orf46, but not with C11orf46Δ. **c** Silver staining of proteins coprecipitated with affinity-purified wildtype C11orf46 (clone 10) or C11orf46Δ (clone 8). SETDB1, MCAF1 and KAP1 were detected in co-precipitate from full-length C11orf46, but not of C11orf46Δ. **d** Top scoring proteins co-precipitated with C11orf46 in cell lines overexpressed with C11orf46 (clone 10), C11orf46Δ (clone 8), and control cells are shown with % of amino acid sequence coverage, see **Supplementary Table 1** for details. **e** Protein abundance (in femtomoles) in affinity-purified C11orf46 complexes. Notice prominence of SETDB1-MCAF1-KAP1 repressor proteins (black). Bar graphs indicate mean±S.E.M. *n* = 3 independent mass spectrometry quantification. **f** Schematic diagrams indicating position of C11orf46 arginine to histidine substitution at codon 236 (R236H) in highly conserved cysteine rich domain (CRD; amino acids 209-237) between human and mouse. **g, h** Co-immunoprecipitation experiments using protein lysates from HEK293 cells overexpressing Flag-tagged C11orf46, C11orf46Δ, or C11orf46^R236H^ showed that absence of CRD domain and R236H mutation is associated with massively weakened or non-detectable C11orf46-SETDB1 binding and partial loss of binding to other components of the complex. Notice that R236H mutation does not fully disrupt C11orf46-KAP1 binding. **i** R236H mutation and structural deletion of CRD at the C-terminal of C11orf46 do not affect binding to histone H3. **j** Flow diagrams for immunoaffinity purification of SETDB1 followed by mass spectrometric analysis. **k** Inducible SETDB1 expression system in HEK293 cell lines (48hours after Dox treatment). **l, m** Silver staining of proteins co-precipitated with affinity-purified SETDB1. Both endogenous C11orf46 and SETDB1 binding partners such as MCAF1 and KAP1 were detected and verified by mass spectrometry. Top scoring proteins co-precipitated with SETDB1 in cell lines overexpressed with SETDB1 (clone 1) and control cells are shown with % of amino acid sequence coverage. For detail of SETDB1-binding proteins that were identified in these experiments, **Supplementary Table 2**. **n** (left) C11orf46-SETDB1 co-immunoprecipitation in postmortem human cerebral cortex. (right) SETB1 co-immunoprecipitates include C11orf46, HP1γ, and MCAF1 in HeLa nuclear extract. o Schematic summary of above data representing working model of C11orf46-SETDB1 complex in relation to histone H3.

### Suppression of *C11orf46* altered a large number of axonal genes

Having shown that C11orf46 is a neuronal nuclear protein bound to repressive chromatin regulators, we hypothesized that the transcallosal dysconnectivty resulting from *C11orf46* knock-down in fetal mouse cortex and of *C11ORF46* deletions in WAGR patients could reflect dysregulated expression of genes critical for axonal growth and development. In order to obtain first insights into C11orf46-sensitive gene expression changes in WAGR deletion cases, we analyzed a RNA-seq dataset of lymphoblastoid cell lines obtained from the aforementioned 2 WAGR cases with *C11orf46* haploinsufficiency compared to 2 controls, including one WAGR case with a shorter microdeletion that retained intact *C11orf46* and one healthy control subject with no loss of genes for the entirety of chr. *11p13-14.1* (**Supplementary Fig. 1b**). We identified 239 differentially expressed genes (FDR <0.05, see Methods; **Supplementary Fig. 3** and **Supplementary Table 3**) that include 41 genes which have KAP1 binding sites ^33^ (**Supplementary Table 4**). Notably, among these 41 genes, we found that 17 genes are reportedly involved in axonal development. However, given that the observed gene alterations were likely confounded by deletion of other proximal genes with C11orf46 at WAGR locus, we switched in our next set of experiments to neuron-like NSC34 Tet-On cells expressing C11orf46 shRNA to further examine C11orf46-regulated genes. We found that expression levels of 17 axonal growth and development genes were altered in response to C11orf46 shRNA (**Fig. 3a**). Next, we wanted to confirm C11orf46’s role as a transcriptional regulator during brain development, focusing on the top three candidate genes from our aforementioned NSC-34 studies, as defined by significant expression changes in response to C11orf46 knockdown and with KAP binding sites proximal to transcription start site (TSS) (+/−750b) ^33^. These include *Doublecortin-like kinase 1 (Dclk1)* which is essential for axon tract formation across the anterior commissure in the ventral forebrain and the corpus callosum ^34^, *Sema6a* which encodes a PLXNA4 protein ligand ^35,36^, and *Gap43* which encodes a growth cone protein essential for commissural axon guidance ^37^ (**Supplementary Table 4**). To confirm C11orf46’s role as a transcriptional regulator during brain development, we knocked down C11orf46 by IUE-mediated delivery of C11orf46shRNA together with GFP expression plasmid at E15 and GFP-labeled projection neurons were isolated by Fluorescence-Activated Cell Sorting (FACS) at P0 (**Fig. 3b**). Notably, *Sema6a* was significantly increased in C11orf46 knock-down projection neurons (**Fig. 3c**). These findings, taken together, strongly suggest that C11orf46 regulates neuronal gene expression, including an inhibitory effect on *Sema6a*.

**Fig. 3.**
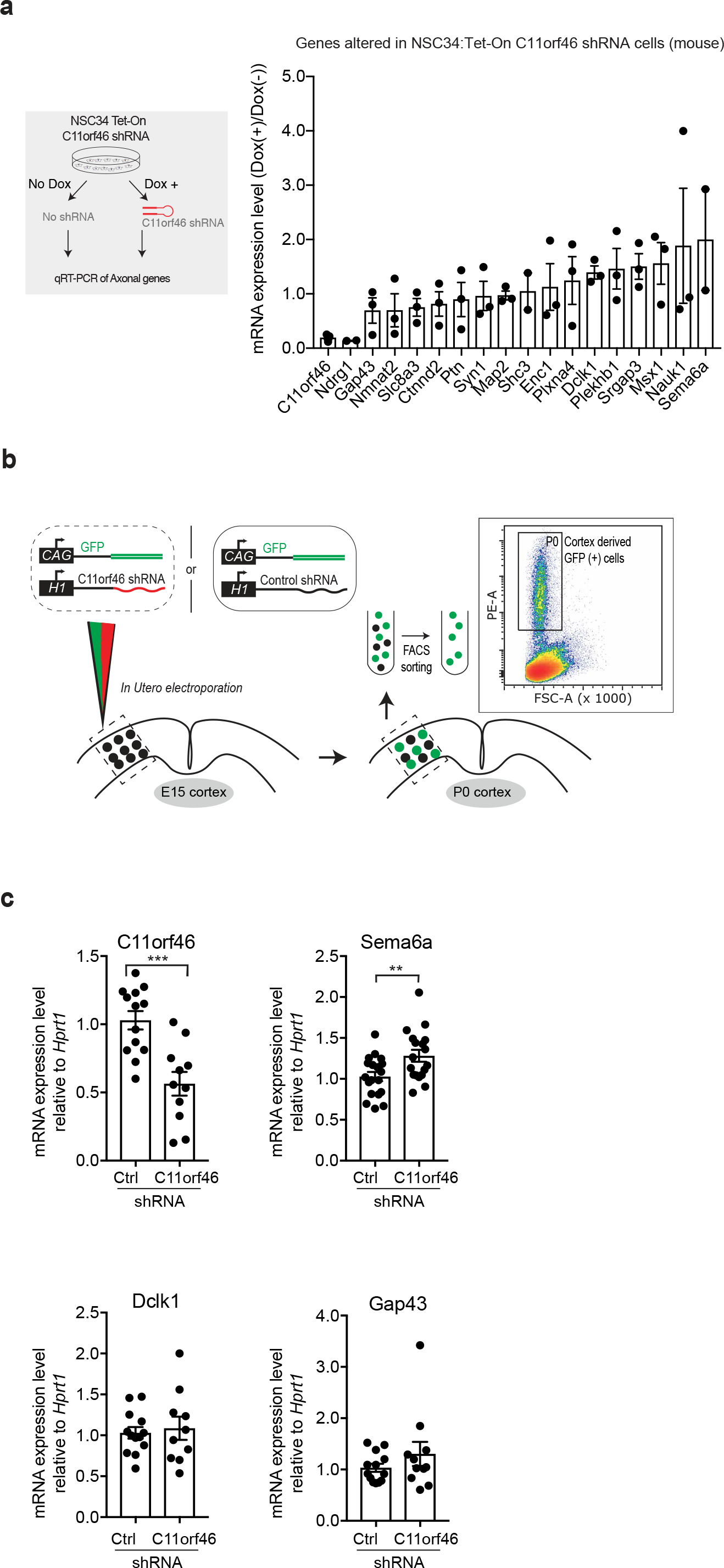
Selection of targeted genes by C11orf46-SETDB1 complex. **a** Mouse neuroblastoma cells NSC34 inducibly expressing C11orf46 shRNA upon doxycycline addition (schematic in left panel); Selected subset of 17 differentially expressed genes implicated in axonal growth and development was tested in RNA expressed from NSC34:Tet-On C11orf46 shRNA cell (right, panel). **b** Neuronal progenitors received control or C11orf46 shRNA and GFP marker plasmids at Somatosensory cortex by IUE at E15; Four days later at P0 GFP positive cells were collected by Fluorescence-Activated Cell Sorting (FACS). **c** *C11orf46* transcript levels were reduced by 50% in sorted GFP-positive cortical neurons electroporated with C11orf46 shRNA1 as compared to GFP-positive neurons with Control shRNA (p=0.0003). Note increase of *Sema6a* transcript upon C11orf46 knockdown (p=0.0085). \**P* < 0.05, \*\**P* < 0.01, \*\*\**P* < 0.001 determined by Student’s t-test test. n = 10-19 independent mice per condition. All data were normalized by the *HPRT1* transcript.

### C11orf46-mediated epigenome editing at neurite-regulating genes rescues a transcallosal dysconnection phenotype

Our results, reported above, showed that C11orf46 co-assembles with the KMT-RC. Apparently, this interaction is important because mutations such as *C11orf46^R236H^*, disrupting C11orf46’s association with the complex, are highly detrimental to orderly brain development. We hypothesized that C11orf46’s affinity to the KMT-RC could be exploited, in order to ‘edit’ neuronal gene expression via a chromatin-associated mechanism. Such type of epigenome editing could be accomplished, for example, by RNA-guided fusion proteins comprised of nuclease-deficient clustered, *regularly interspaced short palindromic repeats* (CRZSPR)/<u>C</u>RISPR-<u>a</u>ssociated protein Cas9 (dCas9) bound to a transcriptional activator or repressor ^38^. However, because ‘1:1’ systems (one transcriptional regulator per dCas9 copy) are often only minimally effective ^38,39^, we instead used dCas9-SunTag protein scaffolds which assembles as a binary system with the protein of interest fused to a single-chain antibody variable fragment (scFv)-green fluorescent protein cassette, with the scFv binding to the Cas9-fused protein scaffold carrying 10 or more copies of GCN4 (the scFv epitope)^39^. We built a dCas9-SunTag system to load ten copies of C11orf46, or the VP64 activator protein as a positive control, onto a single sgRNA-target sequence (**Fig. 4a**). We began with our epigenome editing experiments in HEK293 cells selecting *SEMA6A, DCLK1*, and *GAP43*. For each gene, we obtained four sgRNAs positioned within 500bp upstream of the target TSS. For each of the 3 target genes, three groups of HEK293 cells were compared, based on transfection with (1) dCas9-ST alone (sgRNA-dCas9-10xGCN4^‘SunTag’^) or dCas9-ST together with one of the following plasmids, (2) scFv which recognizes the GCN4 epitope on dCas9-ST-superfold GFP-VP64, or (3) scFv-superfold GFP-wild type C11orf46 (C11orf46^wt^) (**Fig. 4a**). After nuclear localization of the scFv-sfGFP-C11orf46 domain was confirmed (**Fig. 4b**), testing for each target gene experiments were conducted in parallel in HEK293 cells, with each of the four sgRNA tested individually and in a pool of four. Interestingly, introduction of 3 of the 4 sgRNAs to *SEMA6A* which is expressed at much higher levels as compared to the other test genes (**Fig. 4c**), resulting in significant decrease in *SEMA6A* expression when co-transfected with the dCas9-ST^10x C11orf46 (wt)^ systems (**Fig. 4d**). Conversely, *DCLK1* and *GAP43*, which is expressed at extremely low levels at baseline (**Fig. 4d**), showed minimal changes when cotransfected with dCas9-ST^10x C11orf46 (wt)^, while responding to dCas9-ST^10xVP64^ with a robust, at least 5-10 fold increase in their expression (**Fig. 4d**).

**Fig. 4.**
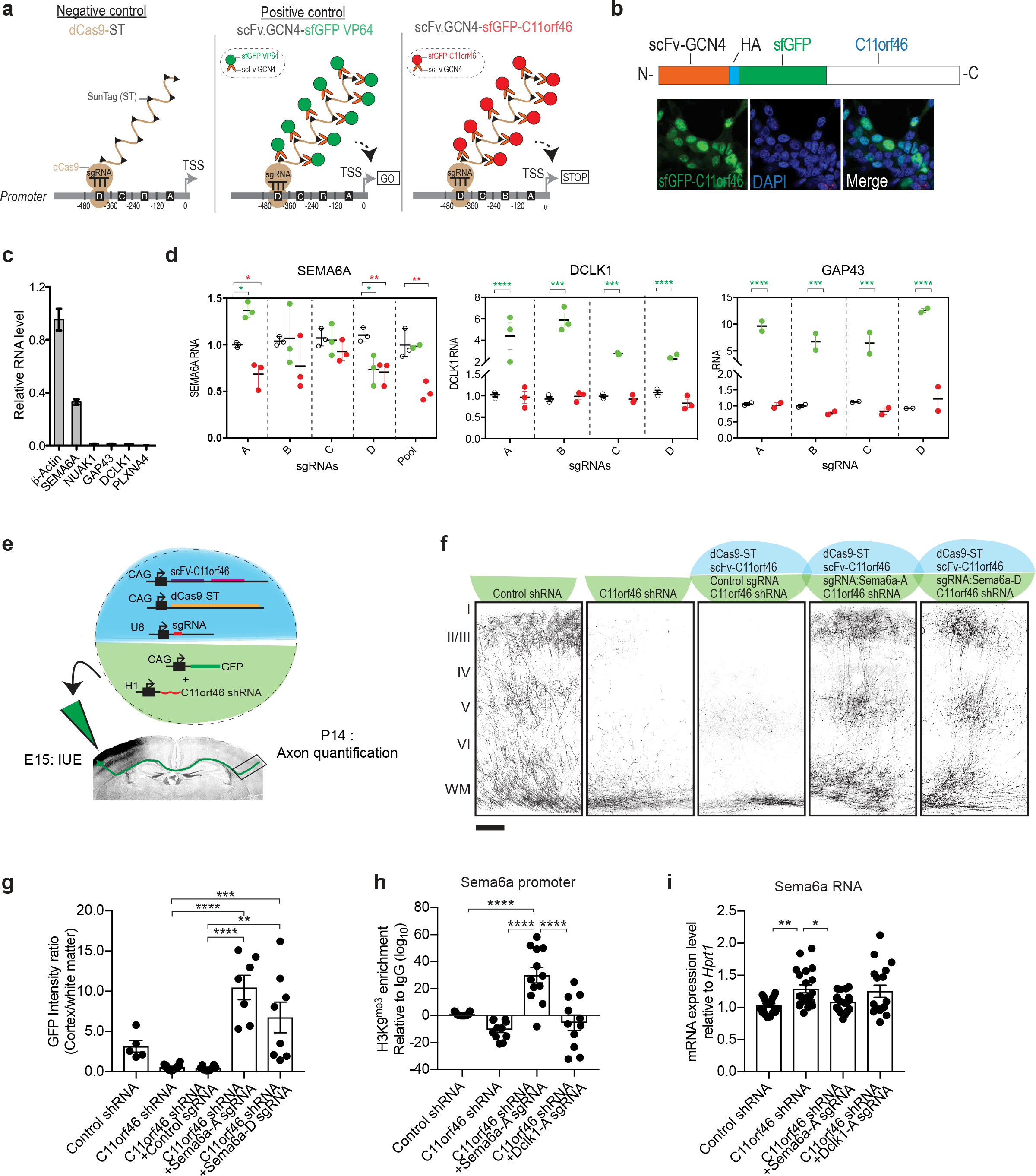
C11orf46-mediated epigenomic promoter editing of neurite-regulating genes rescues transcallosal dysconnectivity in C11orf46-deficient cortical projection neurons. **a** Schematic representation showing epigenome editing approach using nuclease-deficient CRISPR/Cas9 SunTag (dCas9-ST) epigenomic editing system (see text for detail). Sequence-specific single guide RNA (sgRNA) targeting neuronal gene promoters is introduced with dCas9-ST (10xGCN4) and scFv-sfGFP-C11orf46 (or -VP64 as a positive control) in HEK293 cells. **b** Immunocytochemistry of HEK293 with sfGFP-C11orf46 overexpression. fGFP-C11orf46 is well colocalized with DAPI (nuclear) signals. **c** Relative expression levels of neurite-regulating genes in HEK293 cells, notice moderate expression of *SEMA6A*. **d** dCas9-ST mediated promoter loading with 10xC11orf46, and 10xC11orf46^R236H^ in comparison to 10xVP64 at *SEMA6A* in HEK293 cells. For each gene and promoter, 4 sgRNAs (A-D) were tested individually, and for *SEMA6A* also as a pool. Two-way ANOVA followed by Dunnett’s multiple comparisons test was performed where C11orf46 or VP64 mediated epigenetic editing effects were tested by comparing control SunTag system alone. **e** Overview on epigenomic editing strategy of transcallosal projection neurons by IUE. A set of 5 transgenes was delivered simultaneously at E15 by IUE unilaterally into the developing somatosensory cortex, followed by quantification of axonal arborization at P14. **f, g** Axon terminal arborization into the contralateral cortex was disrupted by C11orf46 knockdown, but restored by dCas9-ST mediated recruitment of C11orf46 to two different part of *Sema6a* promoter (Sema6a-), using specific sgRNAs mSema6a-A, mSema6a-D, but not with non-targeting control sgRNA (empty sgRNA cloning plasmid). F (4,36)=13.85165, p<0.0001. \*\**P* < 0.01, \*\**P* < 0.001, \*\*\*\**P* < 0.0001 determined by oneway ANOVA with *post hoc* Bonferroni test. Scale bar, 100μm. **h** H3K9me3 enrichment at *Sema6a* gene promoter by the H3K9me3 antibody in FACS-sorted GFP+ neurons at P0, four days after IUE. Epigenome editing by Sema6a-A sgRNA strongly increase H3K9me3 levels at *Sema6a* promoter. Note that effects are highly target specific, as Dclk1-A sgRNA did not affect H3K9me3 levels at *Sema6a* promoter. F (3,44)=8.718, p=0.0001. \*\**P* < 0.01, \*\*\**P* < 0.001 determined by one-way ANOVA with post hoc Bonferroni test. i Sema6a mRNA level in FACS-sorted GFP+ neurons at P0. Increases of Sema6a expression caused by knockdown of *C11orf46* were suppressed by C11orf46-mediated epigenome editing using Sema6a-A sgRNA, but not by Dclk1-A sgRNA. F (3,72)=4.746, p=0.0045. \**P* < 0.05, \*\**P* < 0.01 determined by one-way ANOVA with post hoc Bonferroni test.

Our studies in HEK293 cells suggest that C11orf46, when tethered in multiple copies to specific promoter sequences via the CRISPR-Cas9-SunTag system, could actively repress transcribed genes. Next, we wanted to explore this approach *in vivo* in developing upper layer cortical projection neurons of somatosensory cortex. We focused on *Sema6a*, which both were de-repressed after C11orf46 knockdown in NSC34 cells and GFP-positive neurons (**Fig. 3a,c**). Two sgRNAs, targeting different sequences within the first 500bp upstream from the TSS of *Sema6a*, were selected for *in vivo* studies and delivered at E15 (**Fig. 4e**). Strikingly these *Sem6a* promoter-targeting sgRNAs fully rescued the transcallosal axonal arborization deficit normally encountered upon C11orf46 knockdown at E15 by IUE (**Fig. 4f,g**). Consistent with C11orf46’s association with KMT-RC, introduction of *Sema6a* sgRNA also resulted in increased H3K9me3 levels at the *Sema6a* promoter, in striking contrast with a decrease in H3K9me3 levels after C11orf46 knockdown (**Fig. 4h**). These effects were highly specific for the target gene sequence, because sgRNAs directed against *Dclk1* promoter did not affect the *Sema6a* promoter (**Fig. 4h**). Likewise, increased *Sema6a* expression in C11orf46 knockdown neurons was strongly suppressed by our SunTag-based C11orf46-mediated epigenome editing toolkit targeting the *Sema6a*, but not the *Dclk1* promoter (**Fig. 4i**)

## Discussion

Axonal development is a pivotal developmental process in establishing neural connectivity for brain functions, including cognition and memory ^40^. This critical stepwise process coordinately regulates the establishment of synaptic connections to target sites with topographic specificity ^41^. Disrupting these processes may lead to altered neural connectivity, which may underlie cognitive disabilities in neurodevelopmental psychiatric conditions such as intellectual disability and autism spectrum disorder ^42^. While multiple molecular pathways including guidance cues and growth factors have been implicated in axonal development ^41^, very little is known about chromatin-regulatory mechanisms ^43^. Recently, the SET (histone methyltransferase) domain-containing transcriptional repressor, PRDM8, was described as essential for the formation of trans-callosal and other interhemispheric fiber tracts ^44^ Likewise, heterochromatin-associated protein HP1γ is required for the crossing of callosal axons into the contralateral hemisphere, while increased neuronal HP1γ expression promotes growth and differentiation of dendrites and axons ^45^. Interestingly, PRDM8 promotes H3K9 methylation ^46^ and HP1γ is bound preferentially to compacted nucleosomal arrays bearing the repressive H3K9me3 mark ^47^ Furthermore, the C11orf46-binding partner and repressive H3K9MTAse, SETDB1, has been implicated in transcriptional regulation of S-type clustered Protocadherin genes, a group of cell adhesion molecules broadly important for neuronal connectivity ^48^. These findings, when taken together with our results reported here, imply H3K9 methyl-regulators as important epigenomic determinant of the developing cortical connectome.

In this study, we showed that C11orf46 assembles with bona fide factors of histone-3 lysine-9 trimethyltransferase (H3K9me3) SETDB1 complex, it substrate histone H3 and a subset of nuclear proteins functionally associated with repressive and heavily H3K9-methylated (hetero) chromatin, including LMNB1, LAP2A and HP1γ. These multi-molecular assemblies of chromatin-bound proteins presumably require distinct domains of C11orf46 to associate with SETDB1 complex and histone H3. Importantly, non-synonymous genetic mutations G218E and R236H found in the CRD domain of C11orf46 are associated with neurodevelopmental disease conditions ^13,49^. We show that specifically the R236H, disrupted the C11orf46 assembly with the SETDB1 complex but not C11orf46’s association with histone H3. Therefore, mutations in critical domains of C11orf46 could disrupt the above assembly and potentially interfere with spatiotemporal gene silencing required for orderly development of transcallosal connectivity. Notably, purified C11orf46 showed strong affinity for histone H3 tail modifications *in vitro;* we were surprised to observe C11orf46’s high affinity towards a histone mark antagonizing histone methylation, H3R2-citrulline (H3R2Cit) ^50,51^. Future studies will explore in more detail the regulatory role for C11orf46. For example, C11orf46 could sense the levels of mono-or di-methylated H3K9 to promote trimethylation, or block H3K9 methylation in response to high levels of antagonizing H3R2Cit mark.

In addition to protein components of the SETDB1 complex, such as KAP1 and MCAF1, our results of proteome screening identified LMNB1 and LAP2A, two nuclear lamina proteins that were previously discovered at the heterochromatin rich nuclear periphery ^52^, as potential protein interactors of C11orf46. Interestingly, LMNB1 and LAP2A play critical roles for developmentally regulated dendritic and axonal connectivity ^53^. Because C11orf46 resulted in impaired axonal development, but had no robust effect on dendritic structure, it will be important to further investigate epigenetic mechanisms to determine how C11orf46 specifically regulates axonal development.

To the best of our knowledge, this is the first study demonstrating the utility of the nuclease-deficient CRISPR/Cas9 system-mediated epigenome editing with IUE knockdown approach for addressing cell-autonomous epigenetic machinery for regulation of transcallosal connectivity. By targeting transcallosal projection neurons, we demonstrate that C11orf46 regulates axonal terminal arborization by control of expression of *Sema6a* via epigenomic promoter H3K9-methylation. The corpus callosum phenotypes in WAGR cases with C11orf46 haploinsufficiency (**Fig. 1**) and the successful epigenomic rescue by sequence-specific C11orf46-mediated chromatin remodeling (**Fig. 4**), would imply that future chromatin-based therapies could be harnessed to correct developmental errors in the brain’s connectome. However, because the WAGR CNV encompasses a large genomic regions that encompass many genes, hampering identification of genetic drivers responsible for specific phenotypes shown in CNVs-associated disease conditions, our data do not exclude the possibility that other genes in 11p13 deletion region, besides *BDNF, PAX6*, and *C11ORF46* may have independent effect on axonal and other anatomical phenotypes as well as behavioral outcomes in WAGR syndrome. Furthermore, since all patients with *C11orf46*+/− also had *PAX6*+/− in our cohort, the phenotype of isolated *C11orf46*+/− is unknown (although homozygous mutations in *C11orf46* have been reported in association with intellectual disability) and there remains the possibility that additional loss of PAX6+/− may be required for C11orf46+/− to cause significant defects in morphologic brain development.

In summary, we identified a chromatin-associated mechanism underlying axonal development and show that C11orf46 plays an important role in transcallosal phenotypes in 11p13 deletion syndrome. Further investigation on epigenetic regulators in CNVs associated with neurodevelopmental disorders and altered neural connectivity is important for translational approaches that pave the way for novel treatment of neurodevelopmental psychiatric conditions in early life.

## Methods

### Human subjects

Patients who had prior genetic testing confirming diagnosis of WAGR/11p13 deletion syndrome or isolated aniridia with known *PAX6* mutation or deletion, as well healthy control subjects who had no chronic medical conditions were recruited through local advertisements and on-line postings. The study was approved by the institutional review board of the Eunice Kennedy Shriver National Institute of Child Health and Human Development, National Institutes of Health, Bethesda, MD and was registered at https://clinicaltrials.gov/ as NCT00758108. Written informed consent was obtained from adult subjects who were competent to provide consent and from the parents or legal guardians of children and adults with cognitive impairment. Clinical studies were performed between December 2008 and May 2014 at the NIH Clinical Research Center.

### Genetic testing

Deletion boundaries for each subject with WAGR syndrome were determined using oligonucleotide array comparative genomic hybridization using a custom-designed microarray platform (Agilent Technologies, Inc., Santa Clara, CA) containing 105,000 60-mer oligonucleotide probes using NCBI Build 36 (hg18) human reference sequence as previously described ^54^ Probes for chromosome 11p were spaced at approximately 400 bp intervals (excluding repeat regions) using 57,925 probes; 121 probes were located within *BDNF*.

### Brain magnetic resonance imaging (MRI)

Brain MRI consisted of one cubic millimeter resolution, T1-weighted images collected on a 3.0 T Philips Achieva MRI scanner with an 8-channel phased array head coil. Corpus callosum volumes were calculated using FreeSurfer’s (version 5.3) subcortical image processing pipeline and published methods ^55^. Briefly, the pipeline uses prior probability of a given tissue class at a specific atlas location, the likelihood of the image intensity given the tissue class, and the probability of the local spatial configuration of labels given the tissue class. The measured corpus callosum volume extends 2.5 mm from the midline on both sides to mitigate against any residual misalignment after registration to the template.

### Lymphoblastoid cell lines

Fasting venous blood samples were obtained from patients (**Supplementary Table 3**). Peripheral blood mononuclear cells were isolated using a Ficoll-Paque gradient (GE Heathcare Life Sciences, Pittsburgh, PA), then infected with Epstein Barr virus, and after sufficient cell line expansion, stored in freezing media at −80°C.

### Antibodies

A mouse monoclonal antibody against Flag-tag (M2) (Sigma, Cat #F1804) was used for affinity purifications. For immunoblots, the following antibodies were used: rabbit monoclonal antibody against the C-terminus of human SETDB1 (Cell Signaling technology, Cat #2196S), KAP1 (cell signaling technology, Cat#4124S); rabbit polyclonal antibody against SETDB1 (Santa Cruz, Cat #sc-66884X), MCAF1 (Novus biological, Cat #NB100-438), Histone H3 C-terminus (Millipore, Cat #07-690). Custom-made mouse monoclonal antibody and guinea pig polyclonal antibody were generated against *E. coli* derived, purified and mass spectrometrically verified full-length human His-tagged C11orf46 protein (**SupplementaryFig. 2b**). For details on the generation of mouse monoclonal antibody, see the references ^56,57^ Altogether, total 45 individually picked anti-C11orf46 mouse monoclonal hybridoma clones were screened by Western blot using *E.coli* derived human C11orf46 as antigen. Five clones (clone numbers 9, 26, 28, 33, and 36) were positive in this assay (clone 26 shown as representative example in **SupplementaryFig. 2b**). Antibodies from these clones were pooled and used for the co-immunoprecipitation assay using tissue homogenates of human prefrontal cortex. The guinea pig antibodies were generated by Cocalico Biologicals (Reamstown, PA) using affinity purified human recombinant His-tagged full-length C11orf46. Rabbit polyclonal antibodies against histone H3 lysine-9 monomethyl (Millipore, Cat #07-450), H3 lysine-9 dimethyl (Upstate, Cat #07-441), and H3 lysine-9 trimethyl (Upstate, Cat #07-442) were used for co-immunoprecipitating endogenous SETDB1 from HeLa nuclear extract.

### Plasmids

Plasmids expressing interfering short hairpin RNA (shRNA) were generated to suppress endogenous C11orf46 protein expression utilizing the pSUPERIOR.puro vector system (Oligoengine). Their target sequences are: C11orf46 shRNA-1 with strong suppression; 5′-CAAACTGAATTTGCTCCAGAA- 3’ and C11orf46 shRNA-2 with milder suppression; 5′- GAAGACAGCTTGTACCTGGTT- 3’. A scrambled sequence that shows no homology to any known messenger RNA was utilized to produce the Control shRNA (5′- ATCTCGCTTGGGCGAGAGTAAG-3′). The HA and scFv-tagged RNAi resistant wild-type human C11orf46 expression constructs containing four silent mutations (underlined) in the target sequence of C11orf46 shRNA (5′- CAAAC<u>A</u>GA<u>G</u>TT<u>C</u>GC<u>A</u>CCAGAA-3′) were produced and cloned into CAG promoter driven plasmid (pCAGGS1vector) for rescue experiments. dCas9-SunTag coding sequence was also transferred into pCAGGS1 vector. Two single guide RNA sequences each *Dclk1* or *Sema6a* promoter region were using online CRISPR design tool^58^. Those protospacer sequences were cloned into sgRNA cloning vector (Addgene #41824) with the protocol distributed on the addgene website (https://media.addgene.org/data/93/40/adf4a4fe-5e77-11e2-9c30-003048dd6500.pdf). The protospacer sequences are listed in **Supplementary Table 5**.

### In vitro epigenomic editing system using dCas9-SunTag-C11orf46

For producing scFv-GCN4-sfGFP-C11orf46, full-length wild type C11orf46 (C11orf46^wt^) or R236H mutant C11orf46 (C11orf46^R236H^) was cloned in-frame to the C-terminus of scFv GCN4 antibody using the BamH1 and NotI sites (addgene plasmid #60904). dCas9-SunTag (2μg, addgene #60903) containing 10 copies GCN4 peptide epitope fused to catalytically inactive Cas9 was delivered into HEK293FT cells (1.5×10^6^, 60mm dish) along with a 4 pooled sgRNAs (50ng each sgRNA, U6-sgRNA-polyT amplicon) to direct GCN4 single-chain antibody conjugated fluorescent C11orf46 (2μg, scFv-GCN4-sfGFP-C11orf46) to neurite-regulating gene promoters. Three days after transfection using lipofectamine 2000, total RNA was extracted and qPCR was performed. scFV-GCN4-sfGFP-VP64 was used as a positive control for activating gene expression. dCas9 alone was tested as a negative control to account background gene activity. For producing U6-sgRNA-polyT, mouse [Dec.2011 (GRCm38/mm10)] and human promoter sequences (500 bp) were identified using PromoterWise (http://www.ebi.ac.uk/Tools/psa/promoterwise/). Four sgRNAs spaced approximately 500bp away from each other in the promoter region of the target genes were identified based on their high gRNA score and low/no off-target site in the genome (**Supplementary Table 5**). sgRNAs were incorporated into U6-sgRNA-polyT cassettes by PCR amplification using U6-Fwd and U6sgRNAter-Rev primers which carried the reverse complement of part of the U6 promoter. These cassettes include the sgRNA (+85) scaffold with guide sequence and seven T nucleotides for transcriptional termination.

### In utero electroporation (IUE)

IUE targeting the somatosensory cortex was performed using our previously published methods with minor modifications ^20,21^. Pregnant C57/BL6 mice were anesthetized at embryonic day 15 (E15) by intraperitoneal administration of a mixed solution of Ketamine HCl (100 mg/kg), Xylazine HCl (7.5 mg/kg), and Buprenorphine HCl (0.05 mg/kg). After the uterine horn was exposed by laparotomy, the shRNA plasmid (1 μg/μl) together with CAG promoter-driven GFP expression plasmids (1μg/μl) (molar ratio, approximately 1:1) were injected into the lateral ventricles with a glass micropipette made from a micropillary tube (Narishige, Cat #GD-1), and electroporated into the ventricular zone of the CP at E15. C11orf46 expression constructs (1 μg/μl) were also co-introduced for rescue experiments. For *in vivo* epigenomic editing, CAG-dCas9-ST (1 μg/μl), scFv-C11orf46 (1 μg/μl) and sgRNAs (1 μg/μl) together with shRNAs and GFP expression plasmid will be introduced at E15. For electroporation, Electrode pulses (40 V; 50 ms) were charged four times at intervals of 950 ms with an electroporator (Nepagene, Cat #CUY21EDIT). All experiments were performed in accordance with the institutional guidelines for animal experiments of Johns Hopkins University.

### RNA-seq and bioinformatic analysis

Total RNA was isolated from four lymphoblastoid cell lines including one derived from healthy control subject, one WAGR patient with *C11orf46 gene* intact and two WAGR patients with *C11orf46* gene deletions using a total RNA extraction kit (Qiagen) according to manufacturer instructions. RNA integrity determined in Bioanalyzer (Agilent’s Technologies) and a total RNA-seq was performed at Eurofins genomics. DNA libraries were prepared from total RNA samples, the adaptor-ligated libraries (125bp) were enriched by PCR amplification and gel purified (average library size of ~270bp). Libraries were sequenced with the Genome Analyzer II (Illumina) using pair-end 50-bp raw fastq reads. The sequencing approach resulted in 32,750,145 and 27,929,697 and 39,292,408, and 54,377,092 reads for lymphoblastoid cell line sample numbers one, two, three, and four, respectively. Sequencing raw reads were
assessed for quality control using FastQC (http://www.bioinformatics.babraham.ac.uk/projects/fastqc/). RNA-seq reads of 50 bp were aligned against the human genome (UCSC hg19) references with TopHat2 using default parameters with Bowtie as the internal aligner and a segment mapping algorithm to discover splice junctions ^59,60^. 77 to 82% percent of all unique sequences correctly aligned to the reference genome. Reads were counted in exon regions using featureCounts tool (subread v1.5.2) producing a table that was normalized to rpkms (**Supplementary Table 3**). Heatmaps were produced scaling rpkm values by scale function (base v3.5.1) and colors were adjusted by quantile breaking to allow colors represent an equal proportion of the data. Two different techniques were used in order to assemble the transcriptome: (i) reference-guided assembly of RNA-seq reads using reference annotations from Refseq ^61^ and (ii) non-reference guided assembly (without the help of a reference gene model) to yield a ‘transcript reference-free’ characterization of RNA-seq reads. For the first approach, BioConductor packages DESeq (References) and EdgeR v2.4.6 in the R programming environment were used. HTSeq-count (http://www-huber.embl.de/users/anders/HTSeq/doc/index.html) was used with UCSC annotation to generate a set of per-gene read counts for each sample, using default parameters including ‘stranded’ set to ‘no’. The HTSeq-count-generated matrix count dataset was used as input. The quartile of genes with the lowest counts was excluded to increase the detection power at a given FDR. For the second approach, the Cufflinks-CuffDiff-CummeRbund pipeline was used ^60^. Fragments per kilobase per millions of reads (FPKM) values were computed for each cell line sample. Genes that were significantly differentially expressed with a FDR of 0.05 in DESeq and EdgeR were selected for further analysis. Both methods yielded highly reproducible results in the expression measurements of the transcripts, and the majority of differentially expressed genes were also significant in an independent analysis using Cufflinks.

### Generation of tet-inducible cell lines

Human full-length *SETDB1* complementary DNA (cDNA), identical to SETDB1 transcript variant 1 (NM_001145415.1; 3876 bp, 1291 amino acids, predicted molecular weight 143 kDa,) was reverse-transcribed using primer Cjk-SA1. Total RNA was isolated from human brain derived U87-MG glioblastoma cells (ATCC, Cat #HTB-14) and was subsequently amplified by PCR (Cjk-SA2 & Cjk-SA4). SalI-Not1 site was double digested and cloned into pENTR4 and transferred to the pFRT TO DESTFLAGHA vector by a LR reaction. The plasmid was completely sequenced. The FLAGHA epitope contributed 17 amino acids to the N-terminal of SETDB1. These amino acids fused at the N-terminus to the ~60bp (1xFLAG)-(1xHA) epitope (FLAG-HA-SETDB1) downstream of a promoter containing the tetracycline-responsive element (pFRT-TO-FLAGHA-SETDB1) and stably transfected (hygromycin) into Flp-In^™^ T-Rex 293™ cells (ThermoFisher, Cat #R780-07) together with plasmid pOG44 (expressing FLP recombinase). Full-length *C11orf46* cDNA (NM_152316.2; 270 amino acids; 783bp) was amplified by PCR from an Incyte plasmid (Openbiosystem, Cat #IHS1380-97432875) using primers Cjk24 and Cjk25, cloned into SalI-NotI restriction sites of the pENTR4 vector. A similar procedure was used to generate HEK293 cells for tet-inducible expression of a FLAG-HA-tagged full length *C11orf46* cDNA (NM_152316.1 784 bp, 260 amino acids). For cloning, high fidelity reverse transcriptase (AccuScript Hi-Fi; Agilent Technologies #600180) and DNA polymerase were used (Pfu Ultra DNA polymerase; Agilent Technologies #600380). All plasmids were sequence verified and the primers used in their generation are listed in **Supplementary Table 6**.

### Affinity purification of protein complexes

Inducible (Tet-On) HEK293 cell lines expressing SETDB1 or C11orf46 or control cells (10^7^ cells/assay) were treated with 2 μg/ml of doxycycline for 48 hours before harvest. Pelleted cells were suspended in CS100-0.02 buffer (Tris HCl, pH 7.4, 10 mM MgCl_2_, 2.5 mM NaCl, 100 mM 0.02% NP-40, 2 units/ml Benzonase, protease inhibitor, and phosphatase inhibitor) at four times, packed cell volume, sonicated, passed five times through a 26-gauge needle, centrifuged at 20,000 x g for 15 min, and filtered through a 0.2-μm filter (Millipore, Cat #SCGP00525). Benzonase was added to eliminate nucleic acid-dependent indirect interactions. Lysate was passed through an anti-FLAG agarose column (packed using 0.5 ml of 50% slurry per 100 mg of extracts) at 4 °C and washed four times with cold CS100-0.02, then four times with cold CS500-0.02 (Tris HCl, pH 7.4, 10 mM MgCl_2_, 2.5 mM NaCl, 500 mM 0.02% NP-40, protease and phosphatase inhibitors). Column-bound protein complexes were eluted using 10 bed volumes of the CS100-0.02-containing 3X-FLAG peptide (100ng/μl) at 4 °C. Purity of protein complexes was verified with four to fifteen percent gradient gel/silver stain and analyzed using mass spectrometry at the proteomics core facility in University of Massachusetts.

### Immunohistochemistry

Immunohistochemistry was performed using our previously published methods with some modifications ^20,21^. Mouse brains were extracted after perfusion with 4% paraformaldehyde (PFA). The fixed brains were embedded in cryocompound (Sakura) after replacement of PFA with 30% sucrose in phosphate buffered saline (PBS). Coronal sections including somatosensory cortex were obtained at 40 μm with a cryostat (Leica, Cat #CM 3050S). The sections were washed with PBS containing 0.5% Triton X-100 and then blocked with 0.5% Triton X-100 and 1% bovine skin gelatin for one hour. After blocking, sections were incubated with mouse monoclonal anti-NeuN (Millipore, Cat #MAB377) and guinea-pig polyclonal anti-C11orf46 primary antibodies at 4°C overnight, followed by incubation with secondary antibodies conjugated to Alexa 488 and 568 (Invitrogen, Cat #A11001 and A11075 respectively) for one hour. Nuclei were labeled with DAPI (Roche, 10236276001). For cell type marker staining, anti-CaMKII (Millipore, Cat #05-532), anti-Olig-2 (Millipore, Cat #AB9610), anti-GFAP (Sigma, Cat #G3893), or anti-Iba-1 (Wako, Cat #019-19741) antibody were used. For analysis of neuronal migration and axonal outgrowth, coronal sections from fixed brains were obtained at 20 μm and 100 μm, respectively. The sections were washed with PBS containing 0.5% Triton X-100 and then blocked with 0.5% Triton X-100 and 5% normal goat serum for one hour. After blocking, sections were incubated with rat monoclonal anti-GFP (Nacalai, Cat #GF90R) antibody at 4°C overnight, followed by incubation with secondary antibodies conjugated to Alexa 488 for one hour.

### Fluorescence-Activated Cell Sorting (FACS)

The mice subjected to IUE at E15 underwent whole brain extraction at P0 or P14. The somatosensory cortex where GFP-positive neurons were localized was dissected using a stereomicroscope with a fluorescent flashlight (Night Sea, Cat #DFP-1). Cells were separated by utilizing papain dissociation kit (Worthington, Cat #LK003150) with minor modification^62^. For ChIP-qPCR experiments, approximate 0.5-1.0 × 10^6^ GFP-positive neurons obtained by pooling 5-6 pups’ GFP-positive cortices were collected by FACS into 1.5ml Protein LoBind Tube (Eppendorf, Cat #Z666505).

### Quantitative real time PCR (qPCR)

FAC-sorted GFP positive somatosensory cortical neurons were suspended into RNAlater solution (ambion Cat#AM7021) and stored at −80°C. Total RNA were prepared and treated by DNAse (PicoPure^™^ RNA Isolation Kit, Applied Biosystems Cat#KIT0204). Total RNA was reverse transcribed to cDNA (SuperScript™ III First-Strand Synthesis System, Invitrogen Cat#18080051). qPCR was performed using PowerUp^™^ SYBR^™^ Green Master Mix (Applied Biosystems Cat#A25742). *Hprt1* was used as a reference gene to normalize the expression data by ddCt method. Target gene primers (Sema6a, Dclk1, Gap43, and Hprt1) are listed in **Supplementary Table 6**.

### Chromatin Immunoprecipitation (ChIP)

FAC-sorted GFP-positive cells were cross-linked with 1% formaldehyde for 15 min at room temperature, quenched with glycine to a final concentration of 0.125 M for another 10 min, and stored at −80 °C overnight before processing. After cell lysates with the lysis buffer (50mM Tris-HCl, 10mM EDTA (pH8.0), 1% SDS, and protease inhibitor cocktail (Roche)), chromatin was sonicated with a Q Sonica Sonicator (45% amplitude, 15second 20 times with 1 minute interval), cleared by centrifugation, and incubated overnight at 4 °C with five to seven μg of the desired anti-H3K9me3 (abcam, Cat #ab8898). Five μg of chromatin was used for each immunoprecipitate. Immunocomplexes were immobilized with 30 μl of protein-G agarose beads (Active Motif) for four hours at 4 °C, followed by stringent washes (wash buffers will be described) and elution. Eluates were reverse cross-linked overnight at 65 °C and deproteinated with proteinase K at 56 °C for 1h. DNA was extracted with phenol chloroform, followed by ethanol precipitation. ChIP-qPCR analyses were performed in a QuantStudio (Applied Biosystems, 12K Flex) using PowerUp™ SYBR™ Green Master Mix (Applied Biosystems, Cat#A25742. ChIP-qPCR signals were calculated as a percentage of input. Fold induction was calculated over a negative genomic region. All primers used in qPCR analyses are shown in **Supplementary Table 6**.

### Quantitative bin analysis of brain slices

The effect of knockdown of C11orf46 on neuronal migration was assessed by performing quantitative bin analysis by our previously published methods ^19–21^. The number of GFP-positive cells in the developing somatosensory cortex, including the cortical plate (CP), intermediate zone (IZ), and subventricular zone (SVZ)/ventricular zone (VZ), were counted. The somatosensory cortex was divided into 10 equal spaces (10 bins) and the percentage of GFP-positive cells in each bin was determined. The numbers of neurons in each category from more than 3 independent experiments were counted in a blinded manner using ImageJ software (http://rsb.info.nih.gov/ij/).

### Dendritic complexity analysis

Dendritic complexity of GFP-positive pyramidal neurons was analyzed by our previously published methods ^21^. Z stacks of images were collected with a confocal microscope (LSM 700, Zeiss, Oberkochen, Germany) using the ×10 objective lens as Z-series of images, taken 9 stacks at 5.0 p,m intervals, 1024 × 1024 pixel resolution at a scan speed of 7 per section. Acquisition parameters were kept the same for all scans. Images were reconstructed by compressing the Z stacks, and then a region of interest was randomly selected (800-μm wide × 400-μm high) in layers I-III of somatosensory cortex where GFP-positive cells were located. Intensity of the GFP signal in the whole area and cell bodies in the region of interest was measured from three images for each brain by ImageJ software. The intensity ratio of GFP fluorescence of the processes (that is, GFP signal in cell bodies subtracted from that in the whole area) to GFP fluorescence in the whole area was analyzed.

### Quantification of the terminal arborization of callosal axons

Axonal arborization was assessed by previously published methods with some modifications ^63^. Tiled scanned images of the somatosensory cortex where GFP-positive transcallosal neurons projected from the IUE-targeted contralateral hemisphere were collected by a confocal microscope (Zeiss, LSM 700 Meta). Images were taken by the 10x objective lens, with averaged four times, and at 0.8-μm intervals with 1024 x 1024 pixel resolution at a scan speed of seven per section. Acquisition parameters were kept constant for all scans. The signal intensity of GFP in the somatosensory cortex was divided by its intensity in the white matter. Five mice were used for quantification for each group. All calculations were performed using ImageJ software.

### Statistical analyses

For analysis of MRI data, ANOVAs accounting for age and sex as covariates compared white matter structure volumes among patient groups. One-way ANOVA with the Bonferroni multiple comparison test was used for factorial analysis among more than three groups. Student’s t-test was used for comparisons between two groups. All data are shown as mean ± SEM, unless otherwise mentioned.

## Supporting information

## Acknowledgements

We thank Dr. Kazuhiro Ishii, Ms. Nicole Rangos, and Mr. Akarsh Sharma for technical and intellectual assistance of experiments; Dr. Xiaolei Zhu, Ms. Aisa Moreno-Megui and Ms. Nada Redradjaja for a critical reading of the manuscript; Lee Blosser and Raffaello Cimbro for sorting cells by FACS;. This work was supported by DA041208 (A.K.), MH091230 (A.K.), AT008547 (A.K.), MH094268 (A.K.), JHU Catalyst Award (A.K.), MH104341 (S.A.), MH117790 (S.A.), the Brain & Behavior Research Foundation (A.K., S.A., A.S.), ZIAHD008898 (J.C.H.), and an NIH Bench-to-Bedside Award (J.C.H).

## Author Contributions

C.J.P., A.S., S.A., and A.K. designed the study. C.J.P. performed biochemical and cellular experiments. A.S. performed biochemistry and *in vivo* experiments. T.F., C.R., A.D., A.G., L.D. assisted biochemical experiments. Y,H., Y.T., E.A. assisted *in utero* electroporation and performed the data analysis of axonal phenotypes and qPCR. G.P. assisted maintaining animals and *in utero* electroporation. J.G.P assisted gene search using Decipher database. S.E. assisted RNA-sequencing data analysis. J.C.H. obtained human samples and generated lymphoblastoid cell lines. J.C. H., F.M.L., J.A.B. obtained and analyzed structural MRI data. C.J.P., A.S., S.A., and A.K. wrote the manuscript. All authors contributed to and have approved the final manuscript.

## Additional information

**Supplementary Information** includes three figures and supplemental tables.

## Competing interests

J.C.H. is the recipient of an unrestricted research grant from Rhythm Pharmaceuticals. The other authors declare no competing interests.

