## Supplementary material for "*In vivo* epigenetic editing of *sema6a* promoter reverses impaired transcallosal connectivity caused by *C11orf46/ARL14EP* neurodevelopmental risk gene"

Supplementary Fig. 1

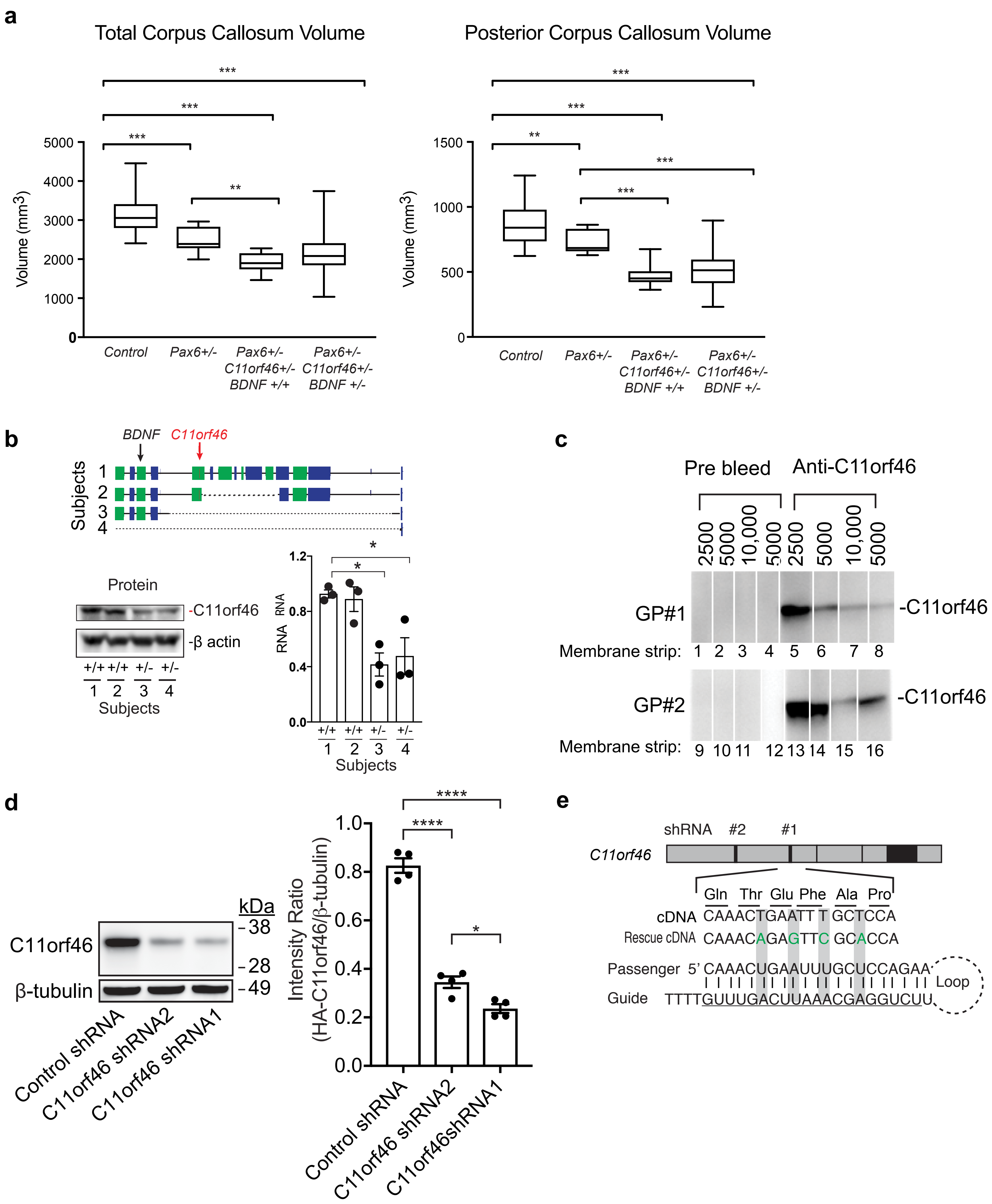

**a**

```
Human   MMDPCSVGVQLRTTNECHKTYYTRHTGFKTLQELSSNDMLLLQLRTGMTL 50
Mouse   -MDPCSVGVQLRTTHDCHKTFYTRHTGFKTLKELSSNDMLLLQLRTGMTL 49
Chicken -MDPCSVGVQLQATNECHKTYYTRHTGFKTKEDISSFDLLLLQLRTGMTL 49
         **********::*::****:*********  :::**  *:***********

Human   SGNNTICFHHVKIYIDRFEDLQKSCCDPFNIHKKLAKKNLHVIDLDDATF 100
Mouse   SGNNTICLHHVKIYIDRFEDLQKSCCDPFNIHKKLAKKNLHVIDLDDATF 99
Chicken SENDTICFHHAKIYIERFEDLQKSCCDPFNMHRKLSKKNLRAIDLHDATF 99
        * *:***:**.*:**:***:**********:*:**:****:.***.****

Human   LSAKFGRQLVPGWKLCPKCTQIINGSVDVDTEDRQKRKPESDGRTAKALR 150
Mouse   LSAKFGRQLVPGWKLCPKCTQIINGSVDVDSDDRQRRKPDSDGRTAKALR 149
Chicken LTAKFGRQFVPGWKLCPKCMQVINGSVDVEAEERQRRKLDSDGRTAKALK 149
        *:******:********** *:*******::::**:**  :*********:

Human   SLQFTNPGRQTEFAPETGKREKRRL-TKNATAGSDRQVIPAKSKVYDSQG 199
Mouse   SLQFTNPGKQTEFAPEGGKREKRKL-TKATSAASDRQIIPAKSKVYDSQG 198
Chicken SLQFTNPGRQTEFTPETSKREKRRLQTKNPSFNSDRQVIPAKSKVYDSQG 199
        ****:***:****:**  .*****:* **  .:  ****:************

Human   LLIFSGMDLCDCLDEDCLGCFYACPACGSTK GAECRCDRKWLYEQIEIE 249
Mouse   LLIFSGMDLCDCLDEDCLGCFYACPTCGSTKCGAECRCDRKWLYEQIEIE 248
Chicken LLLYSGMDLCDCLDEDCLGCFYACPKCGSNKCGTECRCDRKWLYEQIEIE 249
        **::********************* ***.***:****************

Human   GGEIIHNKHAG----------------- 260
Mouse   GGEIIHNKHAGKAYGLLSPCHPYDILQK 276
Chicken GGEIIRNKHVG----------------- 276
         **** :***.*
```

**b**

| # | C11orf46 Peptides Identified by mass spectrometry | Clone-8 (C11orf46 ▲ ) | |
|---|---|---|---|
| 1 | MMDPCSVGVQLRT | + | + |
| 2 | KTLQELSSNDmLLLQLRT | + | + |
| 3 | RTGMTLSGNNTICFHHVKI | + | + |
| 4 | KIYIDRFEDLQKS | + | + |
| 5 | KSCCDPFNIHKKL | + | + |
| 6 | KKNLHVIDLDDATFLSAKF | + | + |
| 7 | TQIINGSVDVDTEDRQK | + | + |
| 8 | RSLQFTNPGRQ | + | + |
| 9 | RQTEFAPETGKRE | + | + |
| 10 | KWLYEQIEIEGGEIIHNKH | - | + |

**c**

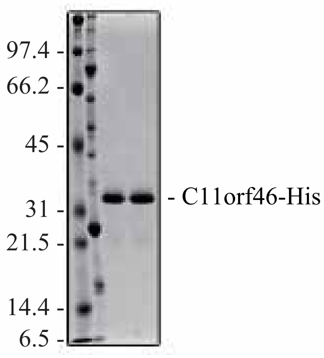


**d**

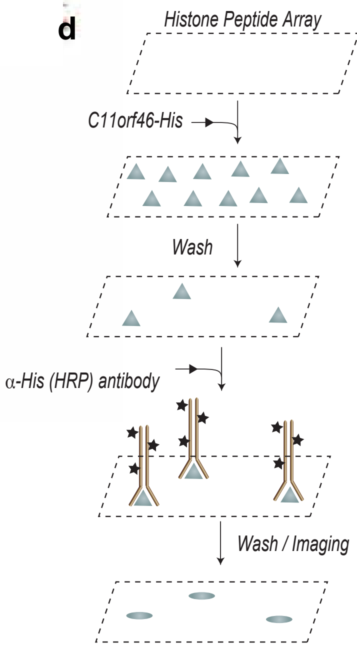


**e**

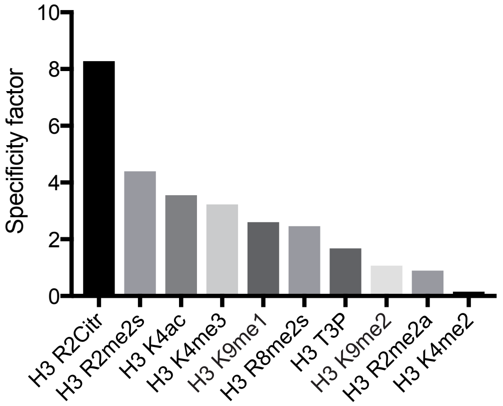


Top 10 (out of 384) modified histone peptides in histone array bound by C11orf46-His

**f**

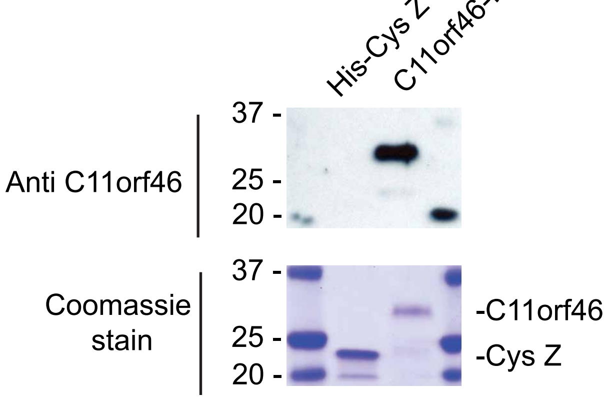

Supplementary Fig. 3

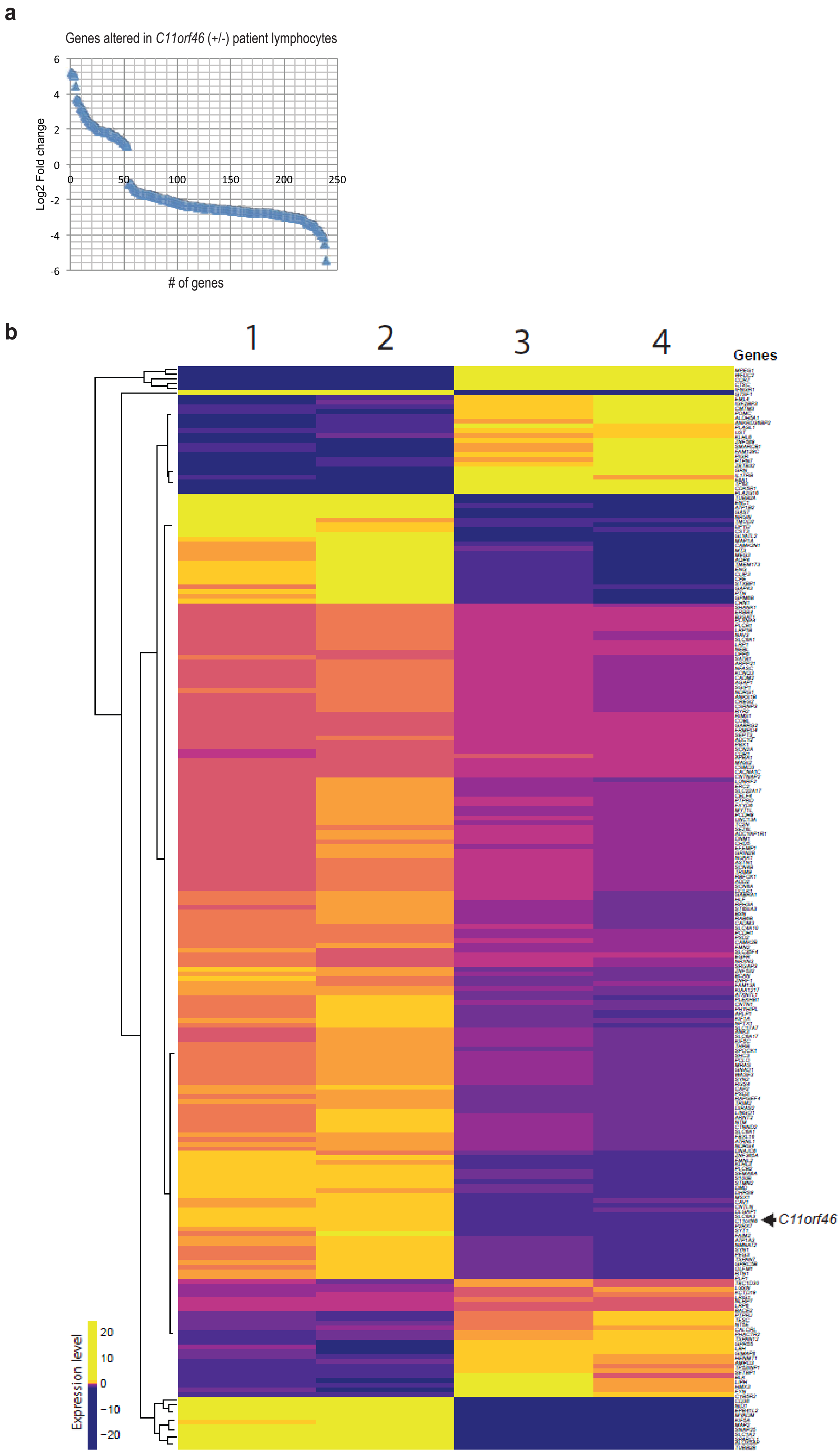

**Supplementary Figure legends**

**Supplementary Fig. 1 The critical role of C11orf46 for interhemispheric connectivity and knockdown effect of C11orf46. a** Total (left) or posterior (right) CC volumes in participants with BDNF and without BDNF. ANCOVA including age and sex as covariates, compared corpus callosum volumes of healthy control (n = 23), isolated *PAX6+/-* (n = 12), *PAX6+/- C11orf46+/-* without *BDNF* haploinsufficiency (n =7) and *PAX6+/-* and *C11orf46+/-* with *BDNF* haploinsufficiency (n =10). No significant differences in the CC volumes of *PAX6+/-C11orf46*+/- patients with and without *BDNF+/-* (p=0.19 and 0.47, for unadjusted comparisons; p=0.13 and 0.37 respectively on ANCOVA adjusting for age and sex). Box and whisker plots represent the distribution of corpus callosum volume from each participant. The inner bar in the box indicates average value. The upper and lower box ends represent first and third quantile, respectively. The upper and lower whisker ends represent the maximum and minimum values in the group, respectively. **b** Schematic illustration of genetic map highlighting various types of micro deletions at 11p13 WAGR locus (dotted lines) including *C11orf46*; one control and three different types of microdeletion in WAGR patients are indicated. Protein (gel image) and mRNA levels of C11orf46 in WAGR patients lymphoblastoid cells were quantified. Bar graphs represent the averages of mRNA expression in three independent experiments per patient. n=3, 1 (*C11orf46* +/+) versus 3 (*C11orf46* +/-) p = 0.0139, 1 (*C11orf46* +/+) versus 4 (*C11orf46* +/-) p = 0.0239. **c** Immunoblot with affinity purified guinea pig (GP) anti-C11orf46 antibodies detecting full-length human C11orf46. Note the pre-immune IgG from same animals (GP#1 & GP#2) did not detect C11orf46 (negative control). Antibody dilutions 1:5000 were used. **d (**Left) Western blot analysis of whole cell extract derived from HEK293 cells cotransfected with HA-C11orf46 and shRNA (C11orf46 or Control) plasmids blotted with anti-HA antibody. Endogenous β-tubulin was used as a loading control. (Right) Exogenous C11orf46 protein expression was strongly suppressed by shRNA1, compared with shRNA2. *$P < 0.05$, ****$P < 0.0001$ determined by one-way ANOVA with post hoc Bonferroni test. **e** Nucleotide sequence targeted by C11orf shRNA1. Four synonimous mutations were added within shRNA1 (bottom) target sequence in the rescue plasmids (green).

**Supplementary Fig. 2 C11orf46 is a chromatin regulator. a, b** C11ORF46 is a conserved molecule. CLUSTAL W multiple sequence alignment of human C11orf46 protein and mouse and chicken *orthologs*. Identical (*), similar (:) critical point mutation associated with cognitive diseases (see text) marked in red. C11orf46 peptides present in C11orf46 complexes purified from clones 8 & 10 are highlighted (bold and underlined) (**a**) and detailed in the table (**b**). Note absence of carboxyl terminal peptide (marked in green) in clone 8 expressing truncated C11orf46. **c** Immunoblot showing recombinant protein load (250ng each). **d** Purified C11orf46-His incubated with Histone peptide array followed by HRP labeled anti-Histidine antibody detection and imaging analysis. **e** Histone peptide array showing purified C11orf46 binding to histone H3 N-terminal tail including its modifications. Top ten histone H3 peptides (out of >300 on the array) detected by C11orf46 includes H3K9me2/3. **f** Immunoblot showing mouse anti-C11orf46 detecting *E. coli* derived histidine-tagged full-length human C11orf46 but not the control protein CysZ; coomassie stained gel (bottom).

**Supplementary Fig. 3 RNA-seq dataset obtained from lymphoblastoid cell lines of the WAGR cases and controls. a** Genes differentially expressed in *C11orf46* haploinsufficient WAGR lymphoblastoid cells. **b** Pair-end (50bp) RNA sequenced libraries were mapped to human (GRCh38.p10_v26) with STAR (v2.5.3a) using a two-method step protocol following tool specifications. Reads were counted in exon regions using featureCounts tool (subread v1.5.2) producing a table that was normalized to rpkms (**Supplementary Table 3**). Heatmaps were produced scaling rpkm values by scale function (base v3.5.1) and colors were adjusted by quantile breaking to allow colors represent an equal proportion of the data.
